# Linear-time prediction of proteome-scale microbial protein interactions

**DOI:** 10.64898/2026.03.01.708874

**Authors:** Andre Cornman, Matt Tranzillo, Nicolo G. Zulaybar, Imane Bouzit, Yunha Hwang

## Abstract

Protein-protein interactions (PPIs) underpin biological function, yet proteome-scale interaction prediction remains bottlenecked by the quadratic computational complexity of all-vs-all pairwise comparisons. Here, we present FlashPPI, a contrastive learning framework, grounded in residue-level interactions, that enables linear-time prediction of physical protein interfaces across a microbial proteome. By leveraging a genomic language model that captures cross-protein co-evolutionary signals from metagenomic sequences, FlashPPI aligns interacting partners in a shared latent space. We demonstrate a four-fold performance increase over existing sequence-based methods, while reducing proteome-wide screening time from days to minutes. Crucially, FlashPPI achieves comparable screening performance to state-of-the-art structure-folding models at a fraction of the computational cost. Finally, we integrate FlashPPI into seqhub.org, an interactive web platform that combines predicted networks with functional annotations and genomic context, making proteome-wide network analysis rapid and accessible for microbial discovery.

## Introduction

The vast majority of sequenced proteins remain functionally uncharacterized [1, 2], a gap that is widening exponentially with metagenomic sequencing [3]. Elucidating protein-protein interactions (PPIs) at the proteome scale offers a promising avenue towards functional annotation of poorly characterized proteins [4] and discovery of novel molecular mechanisms [5]. However, existing computational approaches rely on homology to known interactions [6] or paired multiple sequence alignments (pMSAs) to infer co-evolution [7]. While deep learning has accelerated prediction speed [8, 9, 10, 11, 12], these methods operate on pairs of sequences. Consequently, screening a proteome of *N* proteins requires an all-vs-all comparison with quadratic 𝒪 (*N* ^2^) complexity, rendering large-scale analysis computationally prohibitive (Fig. 1).

**Figure 1.**
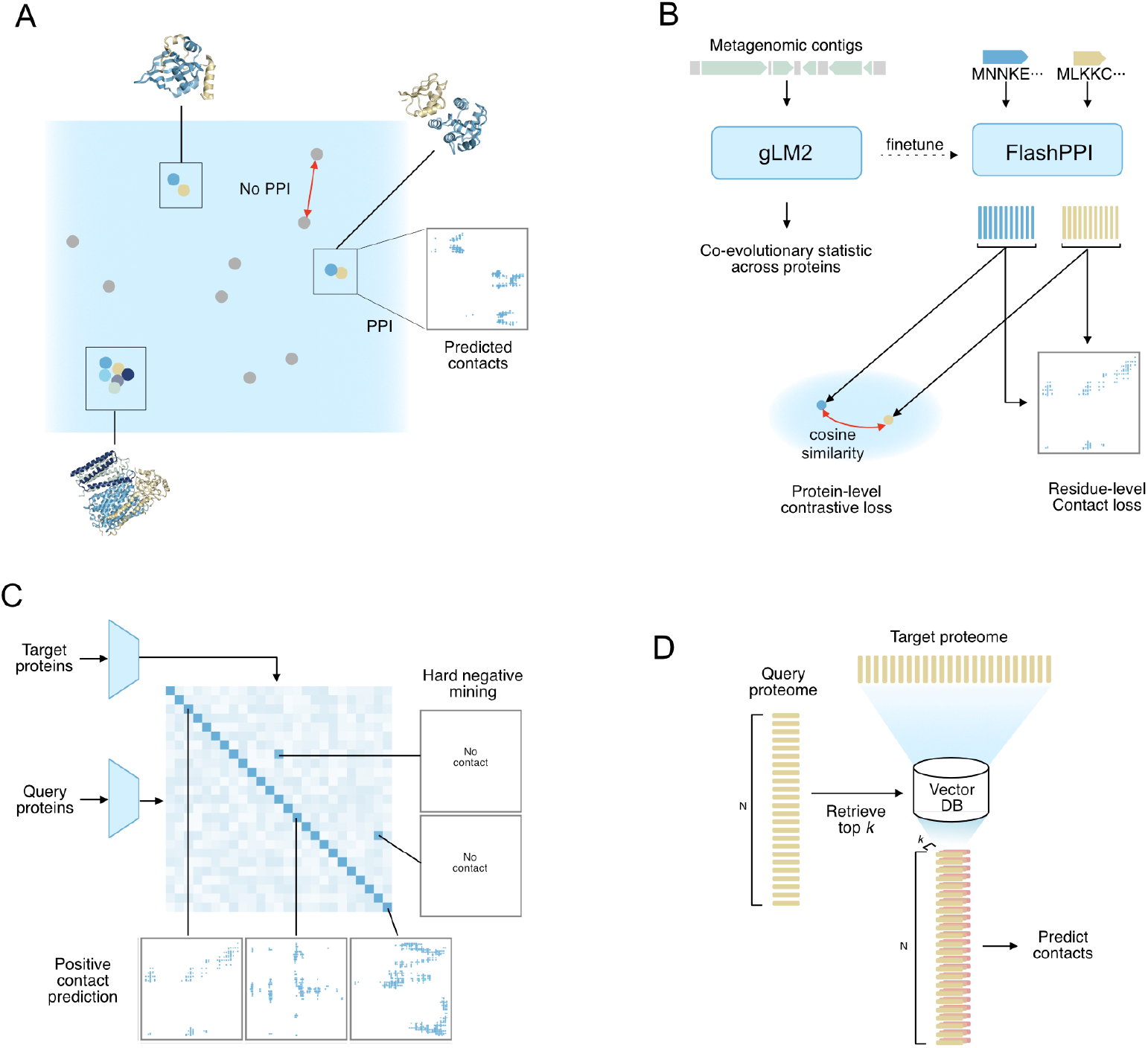
The FlashPPI framework for scalable protein-protein interaction prediction. **(A)** FlashPPI reframes protein interaction prediction as a retrieval task. Proteins are projected into a shared latent space where interacting pairs are close in embedding distance, allowing for the rapid retrieval of candidates that are subsequently verified via fine-grained, residue-level contact map prediction. **(B)** The FlashPPI backbone is initialized with gLM2, leveraging co-evolutionary signals learned from metagenomic contigs. The model is fine-tuned using a joint objective function that combines a protein-level contrastive loss and a residue-level contact loss. **(C)** FlashPPI augments standard in-batch contrastive training with an online hard negative mining strategy. The contrastive similarity matrix is used to dynamically identify hard negatives (non-interacting pairs with high embedding similarity) which are used alongside positive pairs to supervise the contact head and improve discrimination. **(D)** The proteome-scale inference pipeline. The target proteome is encoded into a vector database and the query proteome is used to retrieve the top-*k* nearest neighbors for each query, reducing the search space for contact prediction from quadratic to linear complexity.

We reframe PPI prediction as a dense retrieval task, where interacting proteins are mapped to proximal points in a shared latent space. This formulation reduces the search complexity to linear 𝒪 (*N* ). To bypass the computational bottleneck of generating pMSAs, we leverage the co-evolutionary signals captured by gLM2 [13], a genomic language model trained on multi-protein metagenomic contexts. By combining efficient vector retrieval with fine-grained contact map prediction, FlashPPI achieves both sensitivity and interpretability (Fig. 1). The resulting system processes an entire microbial proteome in minutes on a single GPU (Fig. 2).

**Figure 2.**
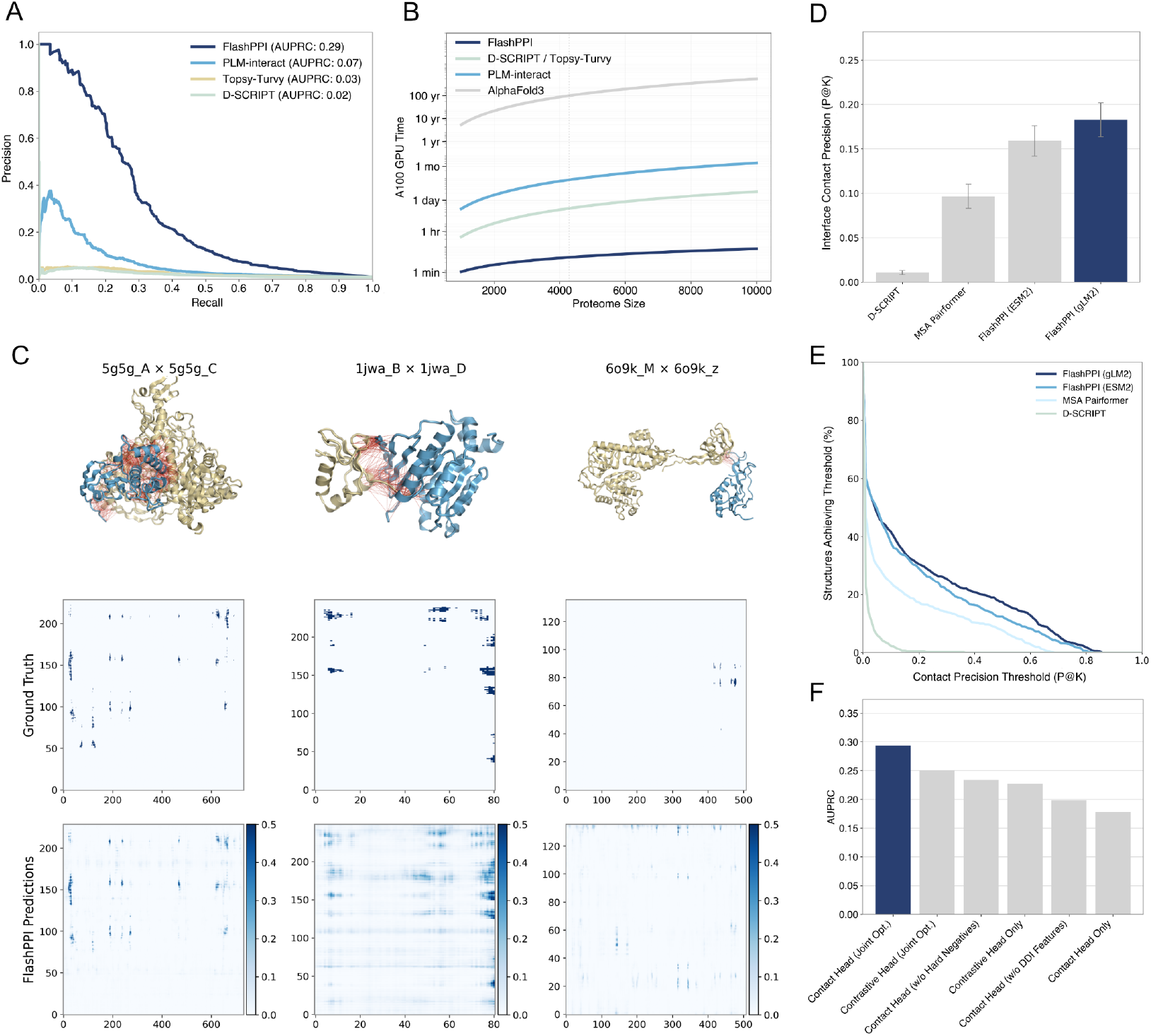
Benchmarking FlashPPI’s predictive performance and speed. **(A)** Precision-recall curves evaluating pairwise PPI prediction on a held-out *E. coli* K12 benchmark (1:100 positive-to-negative ratio). FlashPPI (dark blue) achieves an AUPRC of 0.29, significantly out-performing sequence-based baselines PLM-interact (0.07), Topsy-Turvy (0.03), and D-SCRIPT (0.02). **(B)** Runtime comparison for all-vs-all proteome screening on a single NVIDIA A100 GPU. FlashPPI (dark blue) scales linearly with proteome size, enabling proteome-wide prediction in minutes, whereas baseline methods require days to months for larger proteomes due to quadratic complexity. **(C)** Representative visualizations of predicted contact maps (bottom) versus ground truth (middle) and 3D structures with contacts < 12 Å in red (top) for three *E. coli* test complexes (PDB IDs: 5g5g, 1jwa, 6o9k). FlashPPI accurately recovers interface contacts matching the ground truth. **(D)** Interface contact precision (P@K) on the held-out test set, where *K* is the number of ground-truth contacts. FlashPPI initialized with gLM2 outperforms initialization with ESM2 and baseline methods. **(E)** Distribution of contact precision thresholds achieved across test structures. FlashPPI (gLM2) recovers high-precision contacts for a larger fraction of structures than baselines. **(F)** Ablation study of model components. The full model yields the highest AUPRC (0.29), confirming that hard negative mining and joint optimization are critical for model performance.

### FlashPPI: A sequence-only model for PPI interface prediction

Unlike pairwise classifiers, FlashPPI learns an embedding space where vector distances correlate with interaction likelihood (Fig. 1A). We initialize our model with gLM2 [13], a genomic language model trained on metagenomic sequences that contain multiple (9.7 ± 3.3) proteins per contig (Fig. 1B). gLM2 is a mixed-modality model that represents protein-coding genes in amino acids, while preserving their native genomic relative positions and orientation, as well as any intergenic regions represented in nucleotides. Trained with the masked language modeling objective, gLM2 learns residue-level coevolutionary signals between proteins that are co-located in the genome. Our previous work has shown that this coevolutionary signal can be mapped to known PPI interfaces [13]. We initialize the FlashPPI encoder with gLM2 to leverage this learned cross-protein coevolutionary signal.

FlashPPI employs a dual-encoder architecture with a shared backbone. Protein embeddings are generated via mean pooling and projected by Multi-Layer Perceptrons (MLPs). We optimize the InfoNCE loss [14], maximizing similarity between interacting pairs while minimizing similarity for in-batch negatives. Crucially, we implement false negative masking to prevent penalizing valid interactions that appear as negatives within a batch.

Unlike existing PPI models requiring joint modeling of protein pairs via concatenation [9, 8, 12], FlashPPI encodes protein pairs separately. While this architecture enables efficient interaction candidate retrieval, it precludes the explicit modeling of residue-level interactions across the interface. We recover this structural granularity by co-training a contact head to predict fine-grained interface maps (Fig. 1B). This module, supervised by Protein Data Bank (PDB) contacts (< 12 Å), not only provides interpretability, but is refined via online hard negative mining. We leverage the contrastive embedding space to identify hard negatives during training (Fig. 1C, Materials and Methods M1). This forces the contact head to discriminate true physical interfaces from deceptive non-interacting pairs.

FlashPPI requires a diverse training dataset of physically interacting proteins, paired with ground-truth interface contacts to supervise the contrastive alignment and contact prediction. Accordingly, we utilize the datasets curated by Zhang et al. [11], comprising ∼380k experimental PPIs from the Protein Data Bank (PDB) and ∼530k high-confidence Domain-Domain Interactions (DDIs) from the AlphaFold Database (AFDB) [15]. Because PDB complexes have significant redundancy and bias (e.g., ribosomal subunits), DDI pairs enhance the diversity of the training set and improve model performance (Fig. 2F). To further mitigate training data bias, we cluster PPI sequences at 70% identity and coverage, and employ cluster-weighted sampling, by uniformly drawing training pairs from these clusters.

During inference, the proteome is encoded into a vector database using FlashPPI vector representations. For a query proteome of size *N*, we retrieve the top-*k* nearest neighbors for each protein and predict fine-grained contact maps only for these *N* × *k* candidates (Fig. 1D). This strategy reduces the complexity from 𝒪 (*N* ^2^) to 𝒪 (*N* ), enabling proteome-wide screening in minutes (Materials and Methods M5).

## Results

### Evaluating FlashPPI’s predictive performance

We evaluated FlashPPI on a held-out test set of *E. coli* PPI pairs in PDB dereplicated at 30% sequence identity (*n* = 650). To ensure rigorous generalization, we enforced a strict train-test split by removing any training example where both sequences shared > 30% sequence identity (at > 50% coverage) with the test set. We benchmarked against a challenging 1:100 imbalanced test set, comprising the 650 positive *E. coli* PPI pairs and 65,000 *E. coli* negative pairs. The negative set is composed entirely of proteins known to engage in PPIs (shuffled from the positive set), filtered against STRING (score < 700) to remove accidental true positives. Unlike previous methods that utilize a 1:10 ratio and random negative pairs from the full proteome, this benchmark enforces a realistic class imbalance and utilizes hard negatives composed entirely of proteins known to engage in PPIs.

We compare FlashPPI against state-of-the-art sequence-only PPI prediction methods. FlashPPI achieves a four-fold improvement in Area Under the Precision-Recall Curve (AUPRC) (Fig. 2A) while delivering a 2,400-fold speedup (Fig. 2B) over the next-best model, PLM-Interact [9]. We find that FlashPPI’s contact head outperforms the contrastive head (Fig. S1), and both significantly outperform existing methods. We also find that the contrastively trained embedding similarities correlate well with the contact scores, motivating their use for retrieval of contact prediction candidates (Fig. S2). The multiple orders-of-magnitude acceleration stems from our retrieval architecture, which scales linearly 𝒪(*N* ), whereas pairwise methods like PLM-Interact scale quadratically 𝒪(*N* ^2^).

FlashPPI explicitly predicts residue-level contact maps between retrieved pairs (Fig. 2C). We evaluated contact accuracy on the test set using Precision@*K* (*P* @*K*), where K is the number of ground truth interface contacts [16]. We benchmark our model against D-SCRIPT [8] and MSA-pairformer [17]. D-SCRIPT, while supervised for binary interaction, generates contact maps via unsupervised attention mechanisms. MSA-Pairformer predicts contacts with minimal supervision (*N* = 20 examples), but requires computationally expensive pMSAs as input. In comparison, FlashPPI’s contact prediction is supervised with ground-truth contact interfaces, yielding significantly higher precision on the held-out test set (Fig. 2D, Fig. S3), with a larger fraction of interfaces recovered above varying precision thresholds (Fig. 2E).

We examine the impact of key design choices on model performance. First, we examine the impact of genomic language model initialization. Initialization with gLM2 rather than ESM2 (a protein language model) [18] improves both interaction prediction (Fig. S4) and contact precision (Fig. 2D, 2E), confirming the value of genomic co-evolutionary priors. Next, we investigate the effect of jointly optimizing the contrastive and contact prediction tasks (Fig. 2F). We find that removing the contrastive head (training only on contacts) or removing the contact head (training only on retrieval) degrades performance. Crucially, disabling online hard negative mining, where the contrastive head identifies hard negatives to challenge the contact head, resulted in a significant drop in precision. Finally, ablating Domain-Domain Interaction (DDI) training data reduced performance, highlighting the importance of structural diversity in the training set for generalizing beyond known PPIs.

### Predicting PPI network of the held-out *E. coli* K12 proteome

FlashPPI is designed for proteome-scale interactome prediction via retrieval (Fig. 1A). We next evaluate how FlashPPI generalizes at the proteome level by running the inference pipeline (Fig. 3A) on the held-out *E. coli* K12 proteome. For this inference, we defined high-confidence interactions as those meeting an estimated 70% precision threshold (contact score > 0.61; see Materials and Methods M3). We retrieve the top *k* = 100 matches per protein, as we find that a large fraction (87.5%) of high-confidence pairs are captured within the top 100 candidates (Fig. S5). Next, we remove promiscuous hubs, defined as proteins with > 20 high-confidence interactions (Fig. S6), and filter homologous pairs using BLASTp, as homology often confounds evolutionary-based PPI prediction [11]. Using this pipeline (Fig. 3A), we predict a total of 702 high-confidence interactions across 514 proteins (Fig. 3B and 3C, Table S1). The majority (72%) of these predictions overlap with functional associations in the STRING DB, with 23% matching characterized high-confidence physical interactions (Fig. 3D) with higher FlashPPI contact scores (Fig. S8).

**Figure 3.**
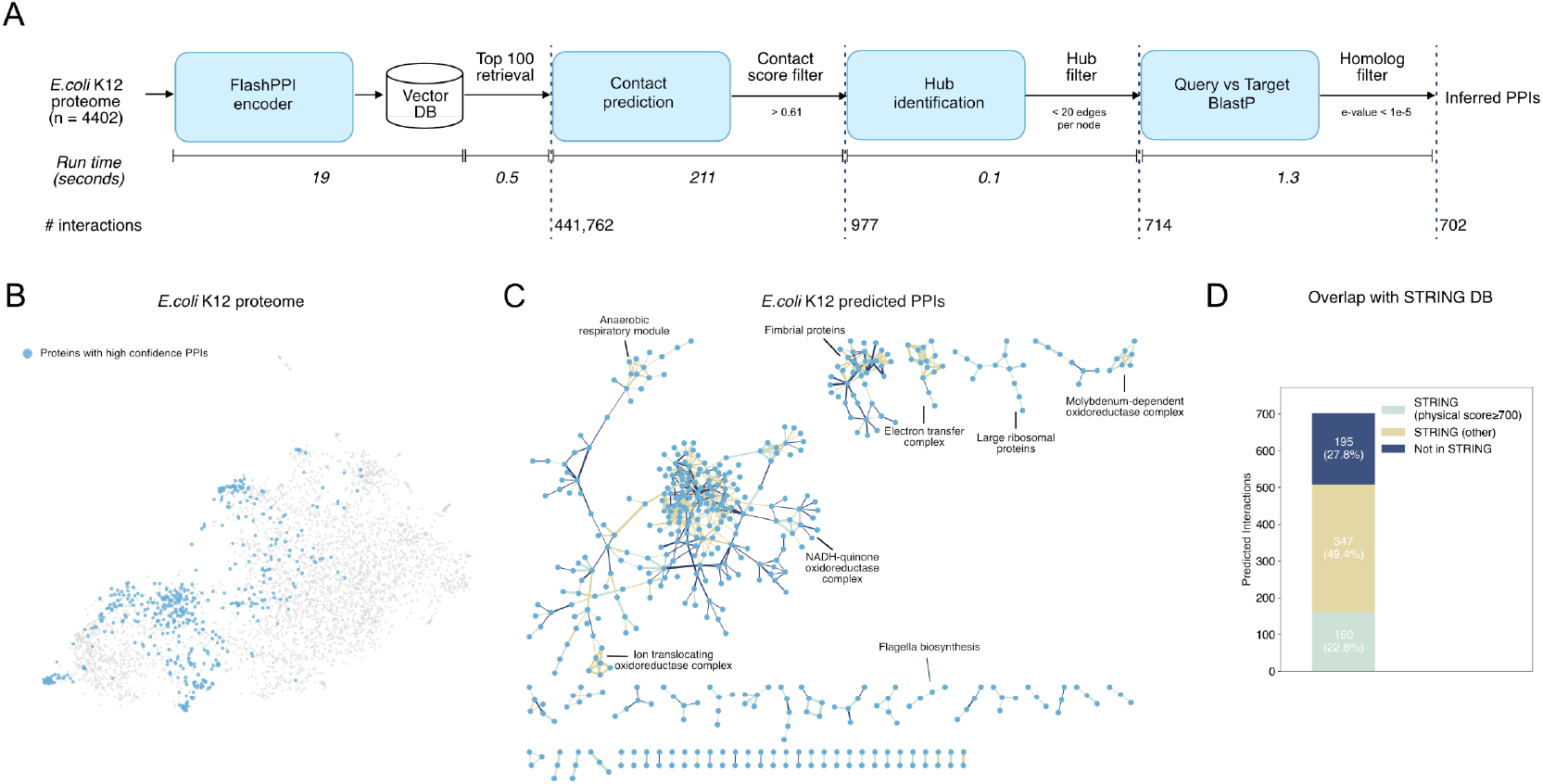
Proteome-scale PPI prediction in *E. coli* K12. **(A)** The end-to-end inference pipeline for the *E. coli* K12 proteome (*N* = 4402). The workflow utilizes dense retrieval (*k* = 100) to propose candidates, followed by fine-grained contact prediction and a cascade of filters: contact confidence (> 0.61), promiscuity (hub filtering), and homolog removal. Runtimes for each stage on a single NVIDIA A100 GPU are shown, demonstrating a total inference time of under five minutes. **(B)** UMAP visualization of the *E. coli* proteome embedding space. Proteins participating in the 702 high-confidence predicted interactions are highlighted in blue. **(C)** The predicted high-confidence physical interaction network. The model recovers distinct functional modules and complexes, including ribosomal subunits, oxidoreductase complexes, and flagella biosynthesis machinery. Edge colors denote whether the interaction has been documented in STRING DB. Navy: not in string, Yellow: STRING (other), Light-blue: STRING (physical). Edge thickness is relative to predicted contact scores. **(D)** Overlap of predicted interactions with the STRING database. The majority of predictions (72.2%) correspond to known associations in STRING (22.8% high-confidence physical, 49.4% other functional), while 27.8% represent potentially novel physical interactions.

To further examine high scoring predictions not documented in STRING, we used AlphaFold2 multimer (AF2-mm) [19] to predict their co-folded structures. We found that AF2-mm also predicts many of these pairs to be interacting, with FlashPPI contact score correlating with AF2-mm size-corrected Interface Predicted Template Modeling (iPTM) scores [20] (Pearson *r* = 0.265, *p*-value = 1.81 × 10^−4^, Fig. S9). Upon manual examination of these putative PPIs, we found that a significant fraction (13.8%) of the pairs were derived from likely false-positive cross-talk between paralogs fimbrial protein and their fimbrial chaperones (Fig. 3 and Fig. S9). We note that discriminating true interacting pairs from cross-talk between sets of paralogs in a diversifying family of proteins (e.g. Fimbrial adhesins and their chaperones [21], toxins and antitoxins [22], PE and PPE proteins [23]) remains a challenging PPI task, even for the latest co-folding models such as AF2-mm (Fig. S9). For such cases, orthogonal evolutionary information, such as genomic co-localization and phylogenetic analysis, can be critical in identifying true positive predictions.

### Comparing FlashPPI to pooled-AlphaFold3 genome-wide screen

AlphaFold co-folding for proteome-wide protein interaction screening is computationally prohibitive and often results in false positive interactions. A recent study demonstrated that pooled AlphaFold3 prediction, where up to 25 proteins are simultaneously co-folded, can lower the false positive rate and reduce inference time per interaction by 2-fold [20]. To compare FlashPPI against AF3-based PPI predictions, we inferred the all-vs-all proteome-scale PPI network of *Mycoplasma genitalium* (proteome size = 476), one of the smallest microbial genomes [24], and benchmarked it against the pooled-AF3 predictions reported by Todor et al. [20] (Fig. 4A). Following their evaluation framework, we use experimentally validated PPIs in STRING DB (referred to as “STRING (experimental)”) as our positive ground-truth set [6]. While both models demonstrate comparable overall predictive performance, FlashPPI achieves higher recall in the high-precision regime (precision > 0.2) (Fig. 4B, Materials and Methods M5). Because global top-scoring prediction lists are frequently dominated by a small number of promiscuous “hub” proteins, we next evaluated the top-*K* interactions on a strictly per-protein basis. This tests the model’s ability to identify the correct partner for any given query. We demonstrate that FlashPPI consistently recovers a larger fraction of known STRING (experimental) interactions among its top-ranked partners per protein than pooled-AF3 (Fig. 4C).

**Figure 4.**
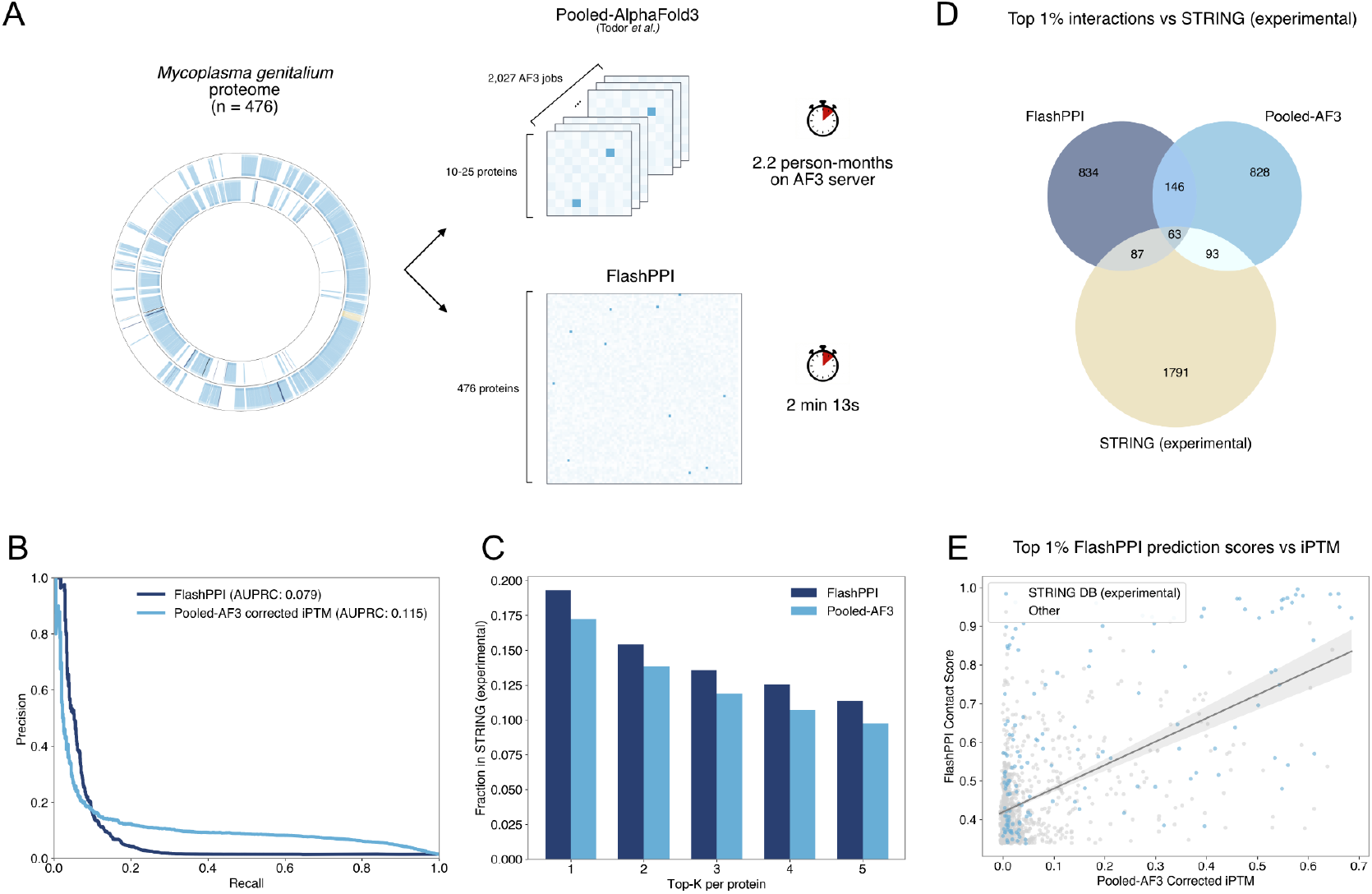
Comparison of FlashPPI to Pooled-AlphaFold3 genome-wide PPI screen. **(A)** Workflow and runtime comparison *M. genitalium* proteome (*n* = 476) PPI screen. FlashPPI screens the entire interactome in ∼2 minutes, bypassing the 2.2 person-months required to prepare and execute 2,027 batched Pooled-AF3 jobs. **(B)** Precision-recall curve evaluating FlashPPI contact scores and pooled-AF3 corrected iPTM scores against experimentally validated STRING interactions. **(C)** Fraction of the top-*K* predicted interaction partners per protein that match known STRING (experimental) interactions, demonstrating higher top-ranked recall for FlashPPI. **(D)** Venn diagram illustrating the overlap of the top 1% highest-scoring pairwise predictions (*n* = 1,130) from each method with STRING experimental data. FlashPPI and pooled-AF3 capture distinct yet complementary subsets of the known interactome. **(E)** Scatter plot showing a statistically significant positive correlation between the top 1% of FlashPPI predictions and their corresponding pooled-AF3 corrected iPTM scores.

We then compared the global top 1% of pairwise interactions (*n* = 1,130) predicted by each method. We found a 17% overlap between the models’ top predictions, with both tools capturing distinct yet comparable fractions of the STRING (experimental) positive set (Fig. 4D). For the top 1% FlashPPI (Fig. 4E) and pooled-AF3 (Fig. S10A) predictions, we observe a statistically significant correlation between FlashPPI contact scores and pooled-AF3 iPTM scores. However, across all possible genome-wide interactions, we find little correlation between the two methods, as many experimentally validated interactions fall into the low-scoring ranges of both models (Fig. S10B). Ultimately, FlashPPI achieves comparable, complementary structural screening capabilities to pooled-AF3 with a ∼20,000-fold faster runtime.

### Cross-proteome PPI in host-virus interactions

FlashPPI can predict cross-proteome PPIs when query and target proteomes differ (Fig. 1D). We explore predicted interactions between previously characterized host-virus genome pairs [25]. We focus our analysis on potentially novel interactions between viral and host proteins, where the query database consists of the viral proteome and the target database consists of the combined set of viral and host proteomes. We identify high-scoring and previously uncharacterized interactions where the viral protein’s highest-ranking match is a host protein (Fig. 5A).

**Figure 5.**
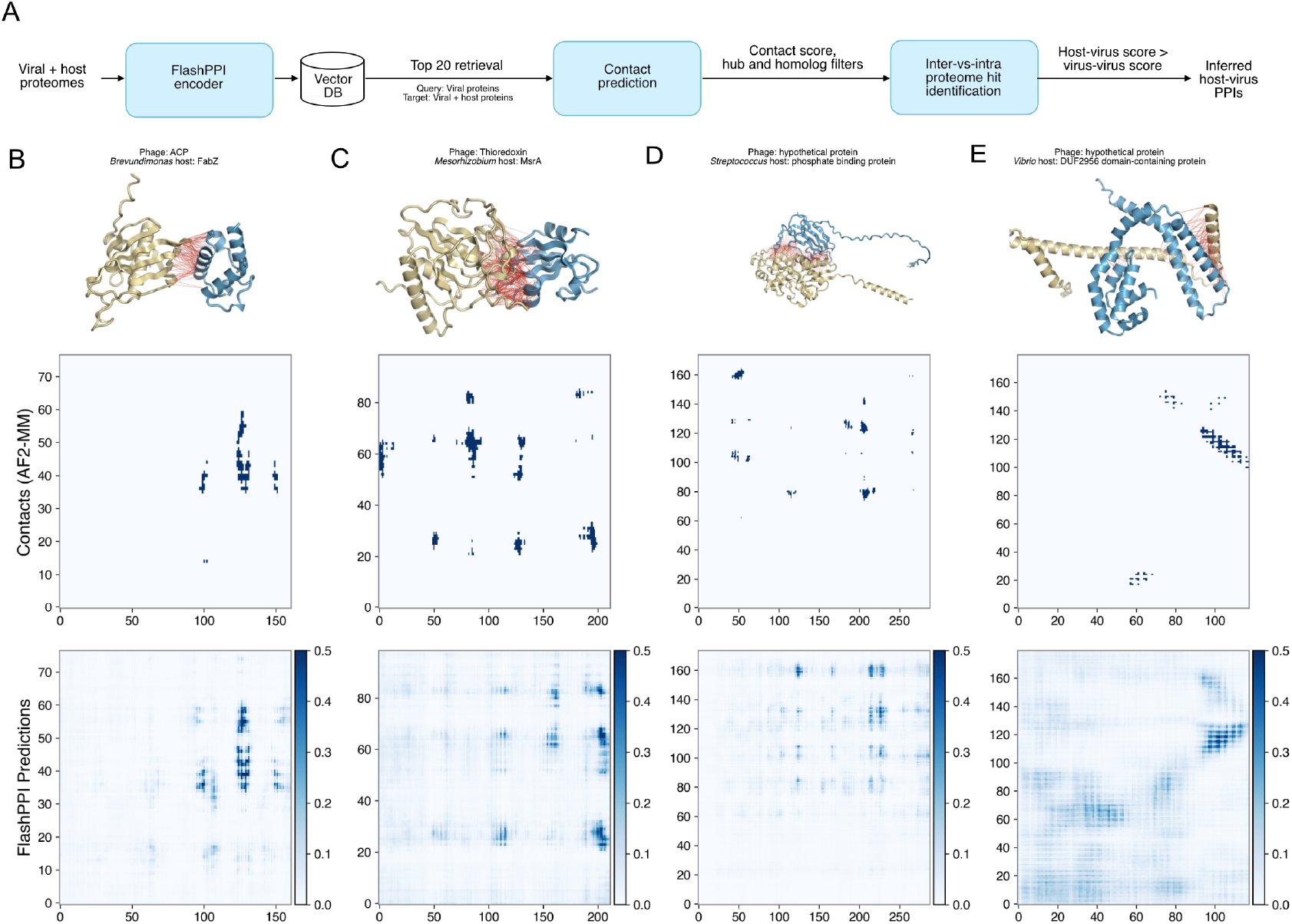
Cross-proteome discovery of host-virus protein-protein interactions. **(A)** The inference pipeline for identifying novel host-virus interactions. Viral and host proteomes are encoded into a vector database. For each viral query, the top-20 viral or host candidates are retrieved. Pairs are filtered by contact confidence, promiscuity (hubs), and homology. High-confidence host-virus interactions are prioritized if the host score exceeds viral-viral interaction scores. **(B to E)** Case studies of predicted host-virus interactions involving poorly characterized proteins. FlashPPI predicted contact maps (bottom) are compared to contacts derived from AF2-MM predicted structures (middle) and 3D complex visualizations with phage proteins in blue, host proteins in yellow, and contacts < 12 Å shown in red (top). **(B)** A phage acyl carrier protein (ACP) predicted to interact with host FabZ, suggesting viral modulation of host lipid metabolism. **(C)** A phage thioredoxin interacting with host Methionine Sulfoxide Reductase A (MsrA), linking viral replication to host redox pathways. **(D)** A hypothetical phage protein targeting a host phosphate-binding protein, potentially enhancing viral replication under phosphate-limited conditions. **(E)** A hypothetical phage protein interacting with a host DUF2596 domain-containing protein, illustrating the ability to recover interactions for conserved proteins of unknown function.

Notably amongst the top-scoring host-virus PPIs, we find previously characterized interactions between phage RNA polymerase sigma factor and host DNA-directed RNA polymerase subunit beta [26], phage ribosomal protein S21 and host ribosomal protein uS11 [27], phage antitoxin and bacterial toxins [28], and phage GroEL chaperonin and host GroES [29] (Table S2). As representative examples, we highlight host–virus PPIs involving poorly characterized proteins (Fig. 5B–E). We predict an interaction between a phage acyl carrier protein (ACP) and host FabZ, suggesting a plausible route for phage-mediated modulation of host lipid metabolism [30] (Fig. 5B). The predicted interaction between a phage thioredoxin and host methionine sulfoxide reductase A (MsrA) points to viral engagement with host redox pathways; thioredoxin-dependent protein-protein interactions are well established in bacteriophage biology, most notably in bacteriophage T7, where thioredoxin is co-opted to support viral DNA replication [31] (Fig. 5C). The predicted association between a phage hypothetical protein and a host phosphate-binding protein is consistent with prior observations that phages frequently target phosphate acquisition systems, including PstS, to enhance replication under phosphate-limited conditions [32, 33] (Fig. 5D). Finally, an interaction between a phage hypothetical protein and a host DUF2596-domain–containing protein is observed (Fig. 5E), illustrating how annotation-independent PPI inference can highlight interactions involving conserved but functionally uncharacterized proteins. Together, these examples illustrate that FlashPPI recovers biologically plausible host-virus PPIs extending beyond canonical transcriptional and translational targets into metabolic, redox, and nutrient acquisition pathways.

### FlashPPI server for fast and interpretable PPI analysis of microbial genomes

To facilitate interactive exploration of proteome-scale interactomes, we integrated the FlashPPI inference pipeline into seqhub.org, a web platform for microbial genomics (Fig. 6). Because FlashPPI’s linear-time complexity circumvents the computational bottlenecks of all-vs-all structural screening, users can generate and visualize whole-proteome networks within minutes by uploading a genome in FASTA format (Fig. 6A). To aid in the discovery of functional modules, the server automatically groups the filtered network into sub-networks using the Louvain community detection algorithm [34] (Fig. 6B). These predicted communities are automatically assigned consensus labels based on the most frequent annotation keywords of their constituent proteins. Predicted PPI communities are mapped directly onto an interactive genome browser, allowing users to cross-reference predicted physical interactions with genomic co-localization (Fig. 6C). This enables the rapid distinction between local (e.g. operonic) complexes and long-range interactions. Selecting a specific interaction edge from a functional sub-network (Fig. 6D) opens a pairwise view that displays FlashPPI’s predicted 2D residue-level contact map (Fig. 6E). This contact matrix is aligned with sequence features (Fig. 6F), including Pfam HMM [35] domains, functional annotations [36], and the genomic distance between the coding sequences. By unifying proteome-wide physical interaction predictions with genomic context and residue-level structural evidence, the FlashPPI server provides a comprehensive environment for accelerating functional discovery in uncharacterized microbial genomes.

**Figure 6.**
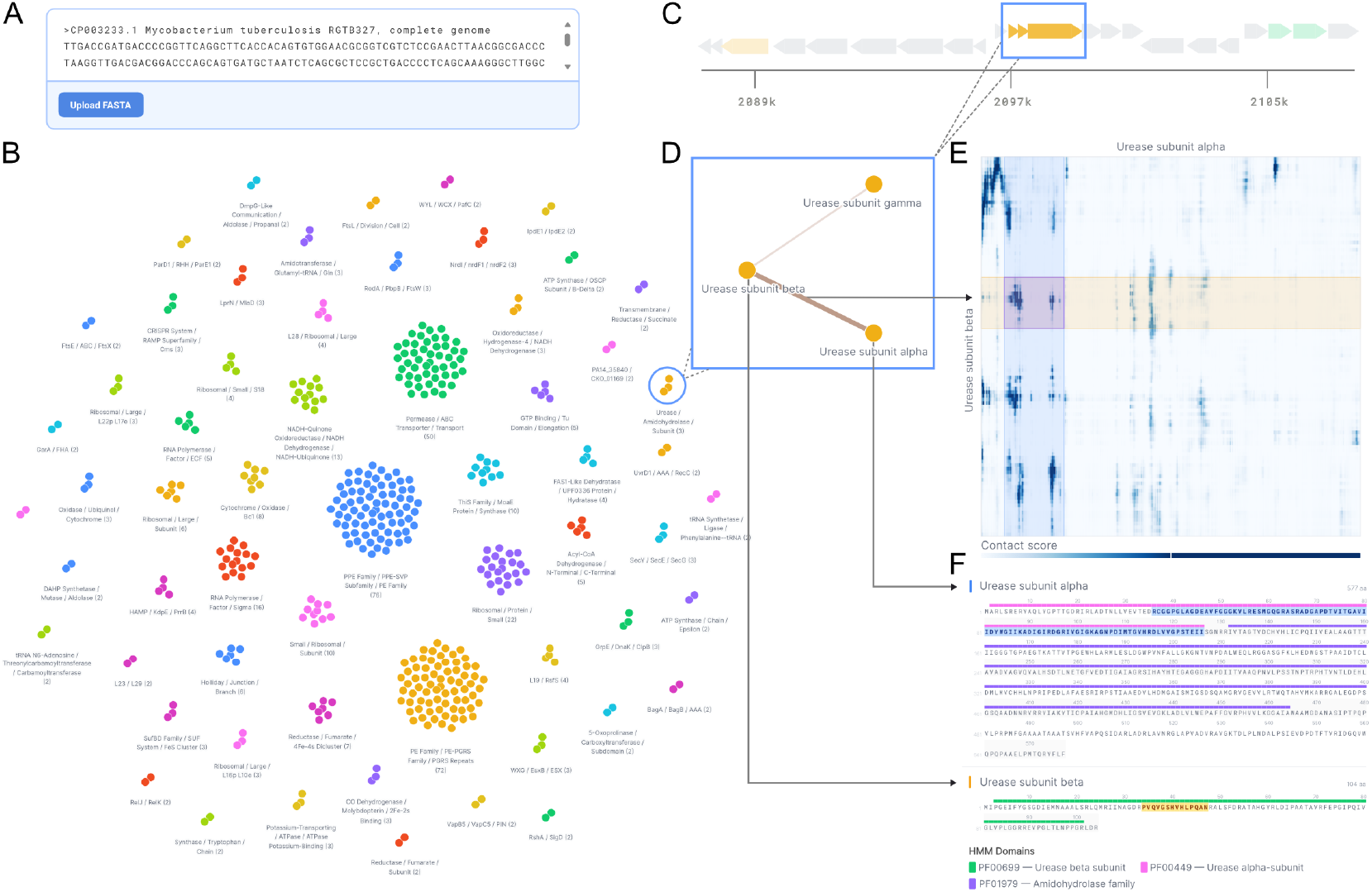
Interactive proteome-scale PPI analysis and visualization on Seqhub.org. **(A)** User interface for initiating proteome-wide predictions via genomic FASTA upload. **(B)** Global visualization of a FlashPPI-predicted proteome network. Proteins are grouped into functional sub-networks using the Louvain community detection algorithm, with consensus labels generated from frequent annotation keywords. **(C)** Predicted interacting proteins mapped onto an interactive genome browser, allowing users to cross-reference physical interactions with genomic co-localization. **(D)** Zoomed-in view of a specific functional sub-network (e.g., Urease complex subunits). **(E)** Interpretable contact prediction displaying FlashPPI’s 2D residue-level contact map for a selected interaction edge. Highlighted regions in the contact map correspond to the subsequence highlights in F. **(F)** Sequence-level visualization aligning the interacting proteins with their functional annotations and Pfam HMM domains.

## Conclusion

FlashPPI reframes proteome-scale protein-protein interaction prediction as a linear-time retrieval task. By coupling global embedding alignment with fine-grained contact prediction, FlashPPI bypasses the quadratic bottleneck of traditional all-vs-all comparisons, reducing screening times from days to minutes while maintaining high precision.

Recent advances in all-atom structural modeling, such as AlphaFold3 [37], provide unprecedented accuracy for multi-protein complexes but remain computationally prohibitive for unguided, genome-scale discovery. FlashPPI bridges this gap by delivering a highly scalable framework for mapping physical interfaces directly from sequence. Crucially, this orders-of-magnitude increase in speed unlocks the systematic exploration of interactomes across diverse, uncharacterized genomes, host-virus ecosystems, and complex metagenomic communities. Integrating fast proteome-wide interaction prediction with targeted high-fidelity structural modeling establishes a powerful paradigm to annotate microbial dark matter and discover functional protein networks across diverse ecosystems.

## Materials and Methods

### M1. FlashPPI Architecture Overview

FlashPPI is a deep learning framework designed to predict protein-protein interactions (PPIs) by aligning interacting pairs in a shared latent space for rapid retrieval, while simultaneously predicting fine-grained physical contacts. The model comprises a protein encoder backbone, query and target contrastive heads for generating dense vector representations, and a contact head for predicting residue-level interactions. The encoder backbone is initialized with gLM2 [13], a transformer-based genomic language model trained on metagenomic data. This initialization leverages gLM2’s learned representations of multi-protein co-evolutionary signals.

#### Contrastive Learning Framework

FlashPPI implements protein interaction prediction as a two-stage retrieval task: vector-based retrieval followed by contact-based re-ranking. We employ a contrastive learning objective to maximize the similarity between interacting protein pairs (positives) while minimizing the similarity between non-interacting pairs (negatives). Given a batch of *N* protein pairs { (*q*_1_, *t*_1_), …, (*q*_*N*_, *t*_*N*_ ) }, we compute the similarity matrix using the dot product of normalized embeddings scaled by a learnable temperature parameter *τ* . The model is trained to identify the correct target for every query within the batch using the InfoNCE objective [14].

#### Residue-Level Contact Prediction and Hard Negative Mining

Fine-grained structural understanding is achieved via a dedicated contact head. Residue-level embeddings from the backbone are concatenated and processed by a 2-layer Transformer module. To predict a 2D interface contact map, we compute multi-head attention scores between each pair of residues, followed by a linear projection layer. The predicted contact map is supervised using binary Focal loss. Following Zhang et al. [11], we define the final contact score as the maximum predicted value in the contact map.

To improve the discrimination of real contact interfaces, FlashPPI employs online hard negative mining. During training, we utilize the contrastive embeddings to identify hard negatives: non-interacting pairs with high embedding cosine similarity. For every protein, we identify the top 2 hard negatives by embedding similarity, and supervise with a contact map of zeros. Additionally, we add a self-negative contact example for each protein to penalize the model for predicting self interactions, which we found to be enriched during early training iterations.

### M2. Training FlashPPI

#### Datasets and Preprocessing

The training data was derived from the PDB training sets curated by Zhang et al. [11]. To ensure rigorous evaluation and prevent data leakage, we removed all sequences sharing > 30% sequence identity at > 50% coverage with the held-out *E. coli* K12 benchmark dataset using MMseqs2 [38]. We further enriched the structural diversity of the training data by integrating the Domain-Domain Interaction (DDI) dataset from Zhang et al. [11], curated from the 200M AFDB [15] structures. As domains are structural and evolutionary units that recombine to create proteins, interfaces between domains often resemble those between distinct protein partners.

The final training corpus consisted of 340,367 full-chain PDB interactions and 529,143 domain-level interactions. To balance the dataset and prevent overfitting to overrepresented protein families, we performed cluster-weighted sampling. We clustered the dataset using MMseqs2 (70% sequence identity, 70% coverage), resulting in 14,729 unique clusters. During training, examples were sampled by first selecting a cluster uniformly at random, then sampling an interaction involving that cluster. DDI examples were sampled with a probability of 0.5 to augment structural diversity.

#### Training Methodology

FlashPPI was implemented in PyTorch and trained on 8 × NVIDIA H200 GPUs. The model was trained for 2 epochs with an effective batch size of 1024 using Flash Attention. To enable large batch sizes, we utilize Gradient Caching [39]. We used the AdamW optimizer with a learning rate of 5 × 10^−4^ and a cosine decay schedule with a 5% linear warmup. To further optimize training memory usage, we froze the first half (16 layers) of the gLM2 backbone during training. The global loss function was a weighted sum: ℒ_*total*_ = 0.1 ℒ_*InfoNCE*_ + 1.0 ℒ_*contact*_. To address false in-batch negatives (randomly sampled negative pairs in a batch that may actually interact) we implemented false negative masking. Using the same 70% identity clustering used for data sampling, we mask any in-batch negative pair (*q*_*i*_, *t*_*j*_) where the protein clusters match the clusters of a known interacting pair. To optimize memory, we employed mixed-precision training (BF16) and gradient checkpointing on the encoder backbone.

### M3. Benchmarking predictive performance on held-out *E. coli* K12 dataset

#### Benchmark Dataset Construction

We evaluate FlashPPI’s ability to generalize to unseen organisms on a held-out benchmark centered on the *E. coli* proteome. Positive interactions were sourced from the PDB, filtered for *E. coli* taxonomy IDs (562, 511145, 83333). We removed redundant pairs using the cluster assignments from Zhang et al. [11] (clustered at 30% identity and 80% coverage). This resulted in 650 unique positive PPI pairs. We generated random non-interacting pairs from the set of positive PPI pairs at a 1:100 positive-to-negative ratio. To ensure the sampled negative pairs are non-interacting, we excluded any pairs listed as physical interactions in the STRING v12.0 database (physical interaction score ≥ 700) [6]. We remove all sequences from the training set that share sequence identity to the benchmark set (see M2).

#### Threshold Calibration

To determine a decision boundary for proteome-scale PPI screening, we create a dataset consisting of the 650 positive *E. coli* PPI benchmark pairs, and randomly sampled negative pairs from the *E. coli* proteome at a 1:100 positive-to-negative ratio, excluding any pairs with documented physical interaction in STRING database. We simulate a realistic PPI ratio of 1:1000 by applying importance sampling weights to the negative class. We computed the precision-recall curve on this weighted distribution and selected the decision threshold that yielded 70% precision (*t* = 0.61). This threshold was subsequently used for all screening analyses.

### M4. Benchmarking interface contact prediction

We assessed the quality of the predicted physical interfaces using the 650 positive complexes from the *E. coli* K12 benchmark dataset. Ground truth contacts were defined as residue pairs with a C*β*–C*β* distance < 12 Å in the experimental PDB structure. For each complex, we ranked all predicted inter-residue contacts by their confidence scores. We reported the Precision at *K* (*P* @*K*), where *K* corresponds to the total number of ground-truth contacts in the interface.

For D-SCRIPT, we extract the unsupervised contact map using `dscript predict –store_cmaps`. For MSA-Pairformer, we extract pMSAs by following their methodology using MMseqs2 proximity-based pairing (distance ≤ 20, coverage ≥ 75%, and sequence identity with query ≥ 30%). We also report *P* @*K* using ground-truth distance < 8 Å, as MSA-Pairformer was supervised with < 8 Å, and find that FlashPPI outperforms existing methods (Fig. S7).

### M5. Runtime estimation and comparison

#### Complexity Analysis

We analyzed the asymptotic computational complexity of FlashPPI compared to state-of-the-art sequence-based methods. Standard deep learning approaches for PPI prediction operate on pairs of sequences, requiring an all-vs-all comparison to screen a proteome of size *N* . This results in quadratic time complexity 𝒪 (*N* ^2^). In contrast, FlashPPI operates in three stages: (1) an 𝒪 (*N* ) linear embedding step where each protein is encoded once, (2) an 𝒪 (*N* log *N* ) (approximate nearest neighbor) retrieval step to propose candidate partners, and (3) an 𝒪 (*N* × *k*) contact prediction step over the top *k* retrieved candidates per query.

In practice we find that the runtime is dominated by model inference (embedding and contact prediction). The 𝒪(*N* log *N* ) approximate nearest-neighbor search contributes a negligible fraction of end-to-end runtime (0.5s for *E. coli*, or 0.2% of total runtime) using FAISS [40], a highly optimized vector search library.

#### Empirical Runtime Measurement

We projected the total runtime required to perform an all-vs-all screen for proteomes of up to 10,000 proteins. For FlashPPI, we measured PPI prediction runtime at retrieval depth of *k* = 100 on 1 Nvidia A100 GPU and found the average embedding time of 3.4 ms per protein and contact prediction time of 0.6 ms per pair. For D-SCRIPT and Topsy-Turvy, we ran both models on a subset of the *E. coli* proteome on 1 Nvidia A100 GPU using the provided CLI (`dscript embed` and `dscript predict`) and measured average embedding time of 102 ms per protein and 4.1 ms per pair for classification. For PLM-interact, we measured average embedding time of 73 ms per concatenated pair following the inference code provided in their GitHub repository (https://github.com/liudan111/PLM-interact). For AlphaFold3, we use the reported latency of 22 seconds per pair (up to 1024 tokens) on 16 A100 GPU (excluding MSA construction time) [37].

### M6. Host-virus interaction prediction

Virus-host genome pairs were obtained from Virus-Host DB [25]. For each virus-host pair, FlashPPI generated embeddings for both proteomes. To comprehensively identify interactions, each viral protein was queried against a combined search space consisting of its own viral proteome and the host proteome. For each viral protein, we search for the top *k* = 20 interaction candidates in the host genome, followed by contact prediction to produce final contact scores. We use *k* = 20 to reduce the computational cost of this analysis, compared to *k* = 100 used for single genome prediction on *E. coli*.

Predictions were filtered for high-confidence physical interactions using a contact score threshold of > 0.4. Interactions involving homologous pairs were filtered using BLASTp (e-value < 10^−5^) as well as promiscuous hub proteins (defined as > 3 interactions). To isolate inter-proteome interactions from intra-proteome interactions, we filtered for viral proteins whose highest-scoring interaction partner was found in the host proteome rather than the viral proteome.

## Acknowledgements

We thank Sergey Ovchinnikov, Joseph Davis, Simon Roux, Michael Laub, and Jeremy Rock for their constructive feedback and ideas on the analysis.

## Funding

This work is funded by the Gordon and Betty Moore Foundation through Grant GBMF13344 to Tatta Bio. Y.H and A.C are supported by Schmidt Futures through Grant G-24-67500 to Tatta Bio.

## Author Contributions

Conceptualization: AC, YH

Methodology: AC, YH

Investigation: AC, YH

Data curation: AC, YH

Validation: AC, IB, YH

Formal analysis: AC, YH

Resource: AC, YH

Software: AC, MT, NZ, YH

Project Administration: AC, YH

Visualization: AC, MT, YH

Funding acquisition: AC, YH

Supervision: AC, YH

Writing-original draft: AC, YH

Writing-review & editing: AC, MT, NZ, IB, YH

## Competing Interests

The authors declare no competing interests.

## Data and code availability

The FlashPPI model is available on HuggingFace at https://huggingface.co/tattabio/flashppi (trained on full dataset) and https://huggingface.co/tattabio/flashppi_noecoli (trained on dataset without leakage to *E. coli*).

Model code and inference scripts are available at https://github.com/tattabio/FlashPPI.

The *E. coli* benchmark dataset is available at https://huggingface.co/datasets/tattabio/ecoli_pdb_benchmark.

All training datasets were downloaded from https://conglab.swmed.edu/humanPPI/humanPPI_download.html.

## 1 Supplementary Material

**Figure S1:**
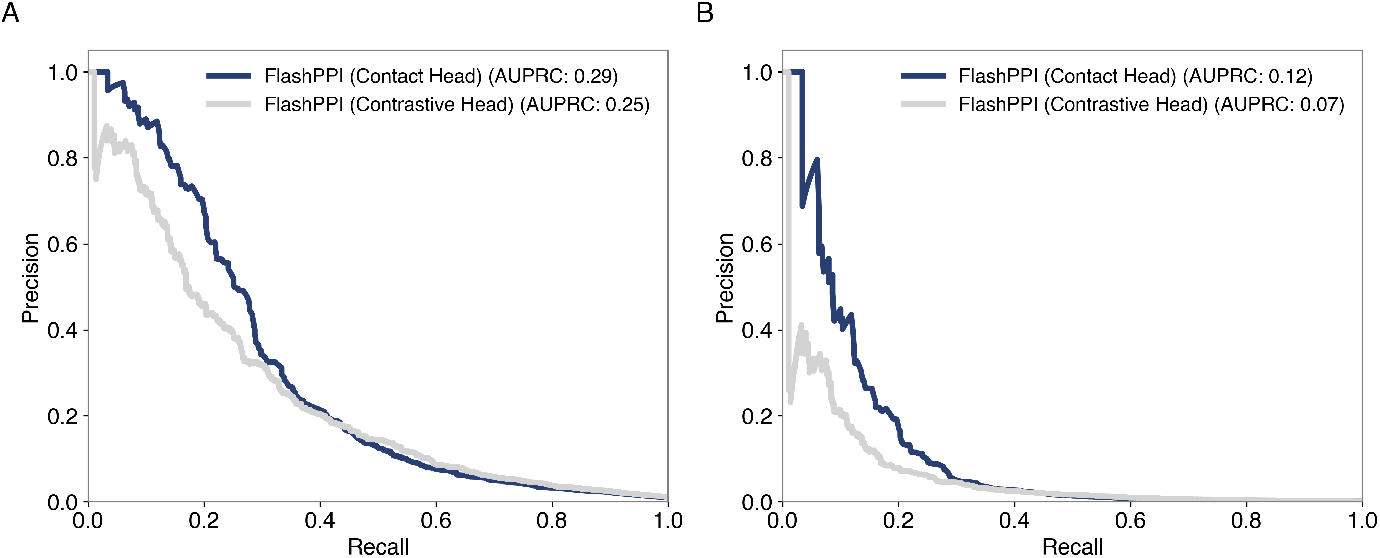
Comparison of classification and contrastive heads. Precision-recall curves comparing the predictive performance of the FlashPPI Contact Head (dark blue) versus the Contrastive Head (grey) on the held-out *E. coli* K12 benchmark at positive-to-negative ratios of (A) 1:100 and (B) weighted 1:1000.

**Figure S2:**
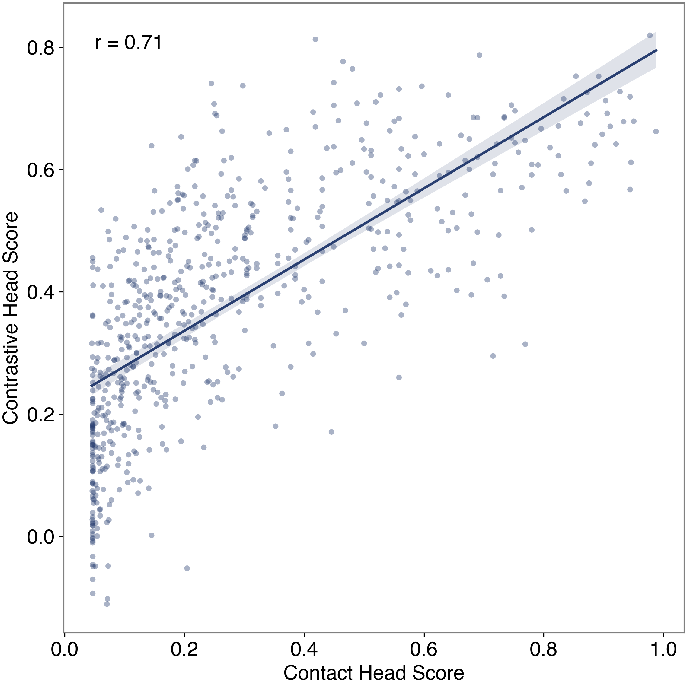
Correlation between embedding similarity and contact scores. Scatter plot showing the Pearson correlation (*r* = 0.71) between the global embedding cosine similarity (contrastive head score) and the maximum predicted contact probability (contact head score) for the *E. coli* benchmark positive pairs.

**Figure S3:**
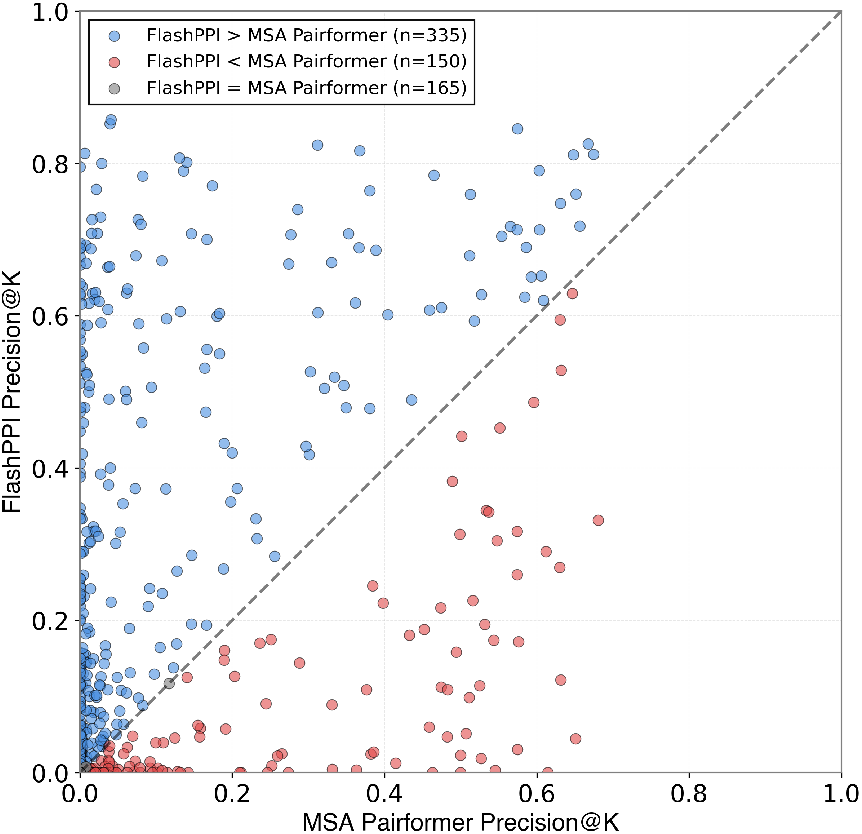
Head-to-head comparison of contact precision. Scatter plot comparing the interface contact precision (*P* @*K*) of FlashPPI versus MSA-Pairformer for each complex in the *E. coli* benchmark set. FlashPPI outperforms MSA-Pairformer on 335 targets (blue), while MSA-Pairformer wins on 165 targets (red).

**Figure S4:**
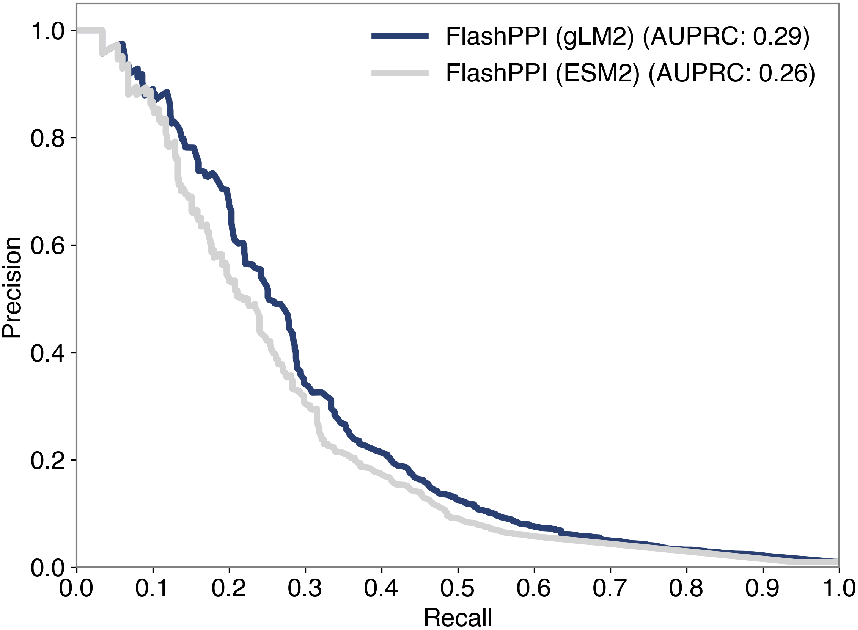
Impact of base model on predictive performance. Precision-recall curves on the *E. coli* benchmark comparing FlashPPI initialized with gLM2 encoder (dark blue, AUPRC=0.29) versus ESM2 encoder (grey, AUPRC=0.26).

**Figure S5:**
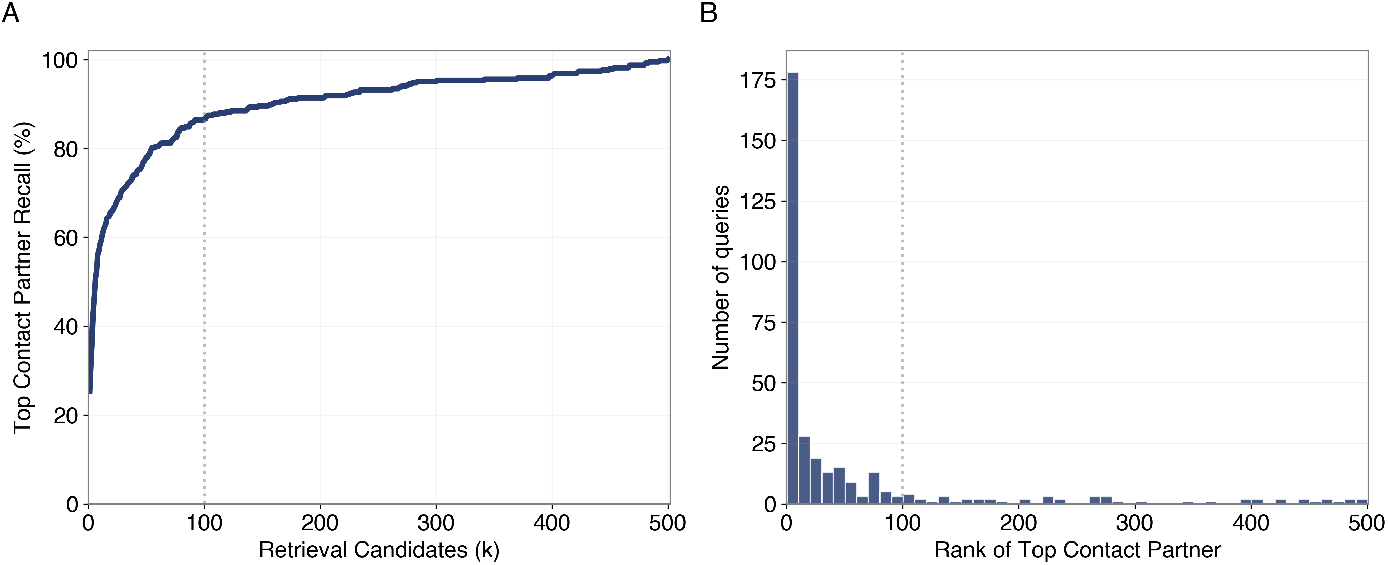
Retrieval efficiency for high-confidence interactions. Analysis of the retrieval depth required to recover predicted physical interactions in the *E. coli* K12 proteome. (A) Retrieval recall for the subset of proteins with a high-confidence interaction (contact score > 0.61). The curve plots the percentage of these proteins where the highest-scoring physical partner is successfully retrieved within the top *k* = 500 candidates. (B) Rank distribution of the highest-scoring contact partner for each query protein. The vertical dotted line indicates *k* = 100; 87.5% of high-confidence interactions are recovered within this window, validating the cutoff used for the proteome-scale inference pipeline.

**Figure S6:**
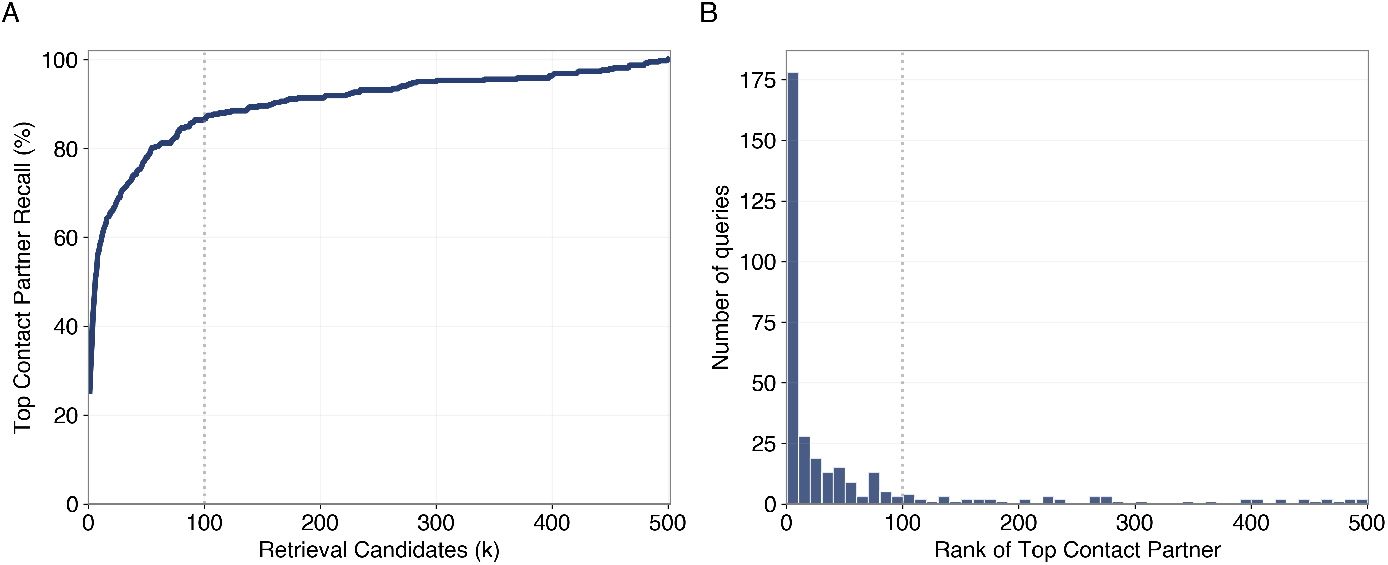
Distribution of predicted interaction degree. Histogram showing the number of predicted high-confidence interaction partners (edges) per protein in the *E. coli* K12 network. The dotted line at 20 edges indicates the threshold used to filter promiscuous hub proteins.

**Figure S7:**
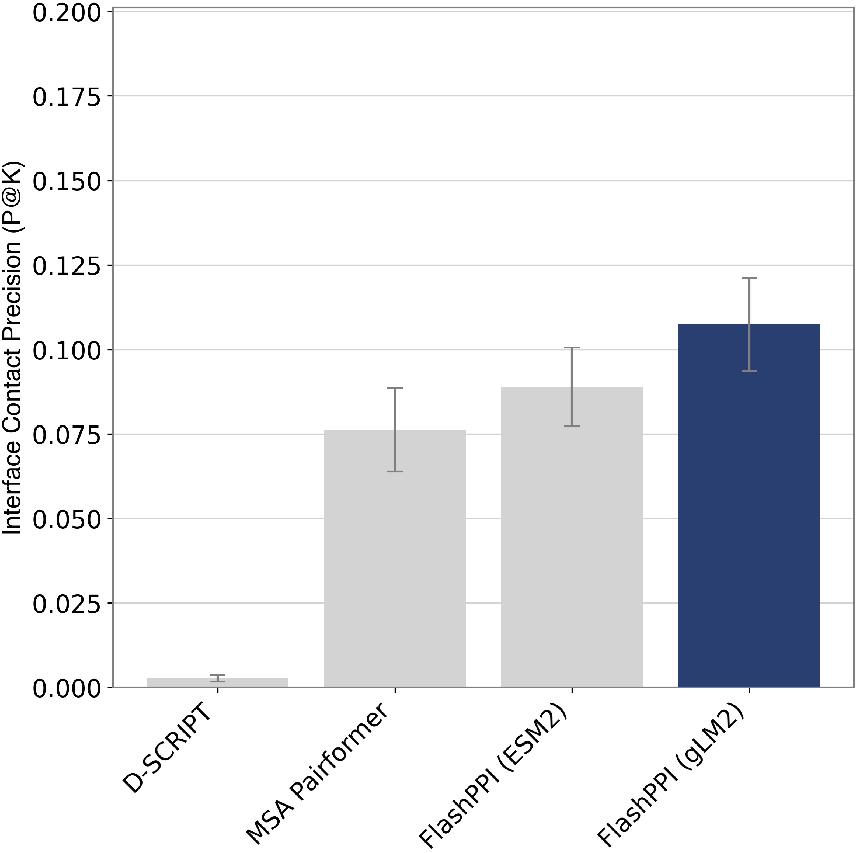
Interface Contact Precision with ground-truth < 8 Å distance. Contact prediction performance comparison evaluated with a ground-truth contact distance of < 8 Å. FlashPPI maintains superior precision over existing methods, including MSA-Pairformer, which was natively supervised using a < 8 Å threshold.

**Figure S8:**
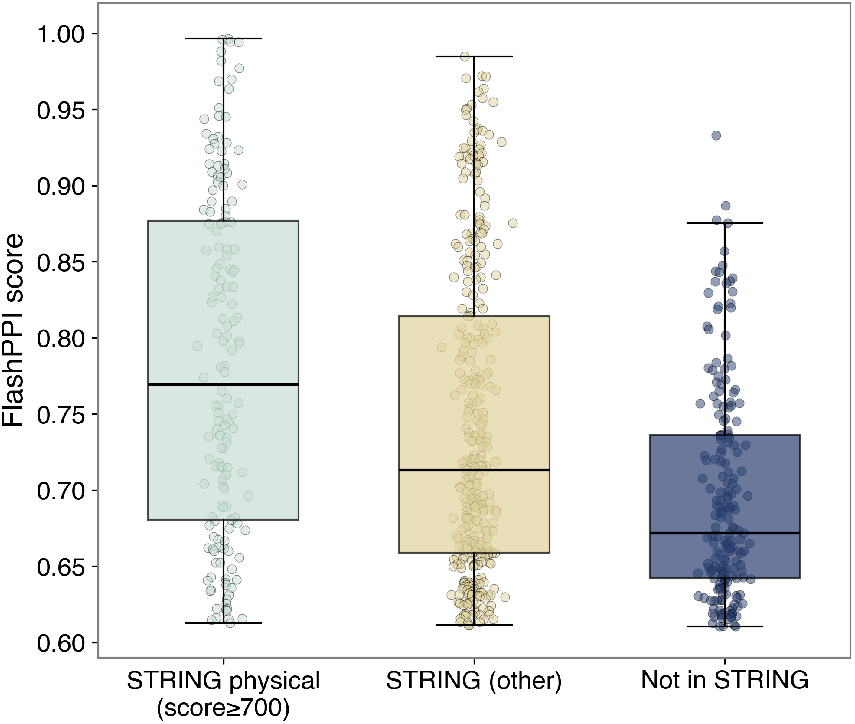
Distribution of FlashPPI scores across STRING characterization categories. Predicted interactions that match characterized high-confidence physical interactions in the STRING database exhibit correspondingly higher FlashPPI contact scores.

**Figure S9:**
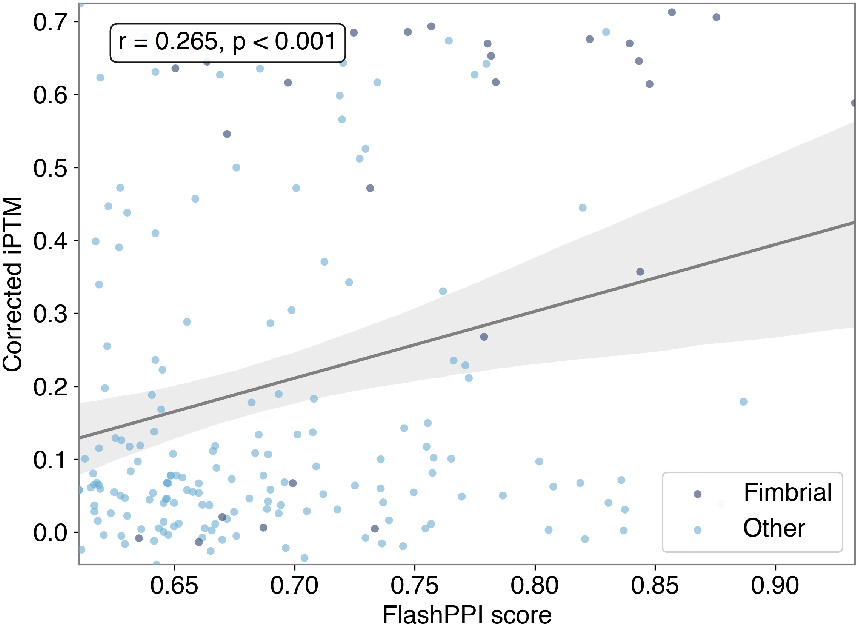
Correlation between FlashPPI and AlphaFold2-multimer (AF2-mm) predictions. Scatter plot demonstrating a weak correlation (Pearson *r* = 0.265, *p*-value = 1.81 × 10^−4^) between FlashPPI contact scores and AF2-mm size-corrected iPTM scores for putative *E. coli* interactions absent from STRING. Dark blue points indicate likely false-positive cross-talk between fimbrial paralogs and their chaperones, illustrating the shared difficulty of distinguishing cognate pairs in diversifying protein families for both models.

**Figure S10:**
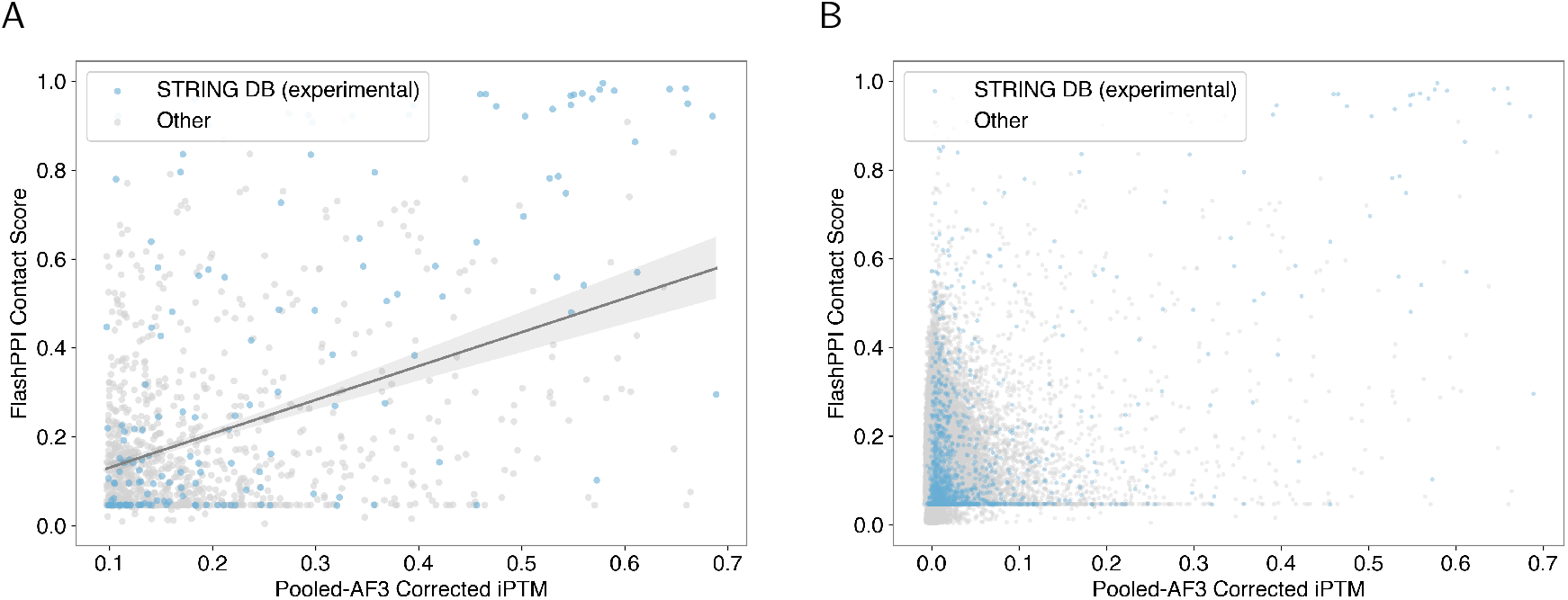
Comparison of FlashPPI scores and Pooled-AF3 Corrected iPTM. (A) Scatter plot of the top 1% of Pooled-AF3 predictions, demonstrating a significant positive correlation (Pearson *r* = 0.429, *p* = 9.6 × 10^−52^) with FlashPPI scores. (B) Scatter plot comparing all pairwise FlashPPI contact scores against pooled-AlphaFold3 corrected iPTM scores. A significant number of STRING interactions fall into lower score ranges for both models.

## Supplementary Tables

- **Table S1:** High-confidence FlashPPI predicted interactions for the *E. coli* K12 proteome (held-out test set).
- **Table S2:** Top-scoring predicted cross-proteome interactions between viral and host proteins.

## Notes

### Competing Interest Statement

The authors have declared no competing interest.

